# Can neural correlates of encoding explain the context dependence of reward-enhanced memory?

**DOI:** 10.1101/2022.09.19.508490

**Authors:** Robin Hellerstedt, Tristan Bekinschtein, Deborah Talmi

## Abstract

Selective encoding can be studied by manipulating how valuable it is for participants to remember specific stimuli, for instance by varying the monetary reward participants receive for recalling a particular stimulus in a subsequent memory test. It would be reasonable for participants to strategically attend more to high-reward items compared to low-reward items in mixed list contexts, but to attend both types of items equally in pure list contexts, where all items are of equal value. Reward-enhanced memory may be driven by automatic dopaminergic interactions between reward circuitry and the hippocampus and thus be insensitive to list context; or it may be driven by meta-cognitive strategies, and thus context-dependent. We contrasted these alternatives by manipulating list composition, and tracked selective encoding through multiple EEG measures of attention and rehearsal. Behavioural results were context-dependent, such that recall of high-reward items was increased only in mixed lists. This result and aspects of the recall dynamics confirm predictions of the eCMR (emotional Context Maintenance and Retrieval) model. The power of ssVEPs was lower for high-reward items regardless of list composition, suggesting decreased visual processing of high-reward stimuli and that ssVEPs may index the modulation of context-to-item associations predicted by eCMR. By contrast, reward modulated the amplitude of Late Positive Potential and Frontal Slow Wave only in mixed lists. Taken together the results provide evidence that reward-enhanced memory is caused by an interplay between strategic processes applied when high- and low-reward items compete for cognitive resources during encoding and context-dependent mechanisms operating during recall.

## Introduction

The ability to selectively encode information that has relevance for future goals is a crucial function of long-term memory. For example, it is more important to remember details of a medical appointment than a casual conversation with a stranger on the bus to the hospital. Selective encoding can be studied in the lab by manipulating how valuable it is for participants to remember specific stimuli, for example by varying the monetary reward participants receive for recalling a particular stimulus in a subsequent memory test. Confirming prevalent intuitions, research shows that people remember information that predicts future reward better than other information (for review see Knowlton & Castel, 2022).

It has been suggested that given our limited encoding capacity, increased memory performance for reward-related information is due to the propensity to engage prefrontal metacognitive encoding strategies to selectively encode valuable information and ignore less important information when both are encountered in the same context (Cohen et al., 2014, 2017). Alternatively, reward-enhanced memory may be driven by automatic dopaminergic interactions between reward circuitry and the hippocampus (Adcock et al., 2006; Bowen et al., 2020; Gruber et al., 2016; Shohamy & Adcock, 2010) and occur regardless of whether high reward and low reward information are encoded in the same or separate contexts. These two views are not mutually exclusive and may operate together (Cohen et al., 2019), but evidence for context dependence of reward-enhanced memory cannot be explained by automatic reward processes alone. The first aim of this study was to investigate if reward-enhanced memory is context-dependent by using a list-composition manipulation.

The list-composition effect refers to an interaction between how items are processed during encoding and the composition of the encoding list (McDaniel & Bugg, 2008). List composition is manipulated by having participants encode two types of items in the same list context (mixed lists) or in separate list contexts (pure lists). It has been shown that emotion-enhanced memory, i.e. increased memory for items with negative valence compared to neutral valence, is dependent on list context. The effect is present when the items are studied together in mixed lists, but is decreased or disappear altogether in pure lists where negative and neutral items do not compete during encoding and retrieval (Barnacle et al., 2016, 2018; Talmi et al., 2007; Talmi & McGarry, 2012). Of relevance for the present study, we recently provided initial evidence that *reward*-enhanced memory also is context dependent (Hellerstedt & Talmi, 2022; Talmi et al., 2021). In the forementioned studies, we found that reward increased memory performance in mixed lists, where high and low reward items were encoded together, but not in pure lists where they were encoded separately. The first, pre-registered hypothesis (link to pre-registration: https://osf.io/cndzs/?view_only=0bc346eef0e9404a82d526b2db40f3fe) is that we will replicate this pattern of results here, using a larger sample and improved materials.

Strategic, meta-cognitive mechanisms could either operate during *encoding* or during *retrieval*. On the one hand, selective attention, selective rehearsal or semanticization (see for example (Cohen et al., 2014, 2017) could be applied during *encoding* to prioritise encoding of high-reward items. On the other hand, the list composition effect could be explained by competition during *retrieval*. One cognitive computational model that formalizes how list-composition effects could occur is the emotional Context Maintenance and Retrieval Model (eCMR, Talmi, Lohnas, et al., 2019) which has successfully modelled list-composition effects both in the context of emotion (Talmi, Lohnas, et al., 2019) and reward (Talmi et al., 2021). Similarly to previous computational models of list-composition effects (Shiffrin & Steyvers, 1997), this model explains emotion and reward-enhanced memory effects as the consequence of an interaction between encoding and retrieval processes. Specifically, in eCMR the prioritisation of high-reward items at encoding only yields an advantage in a free-recall test when these items compete with low-reward items for retrieval. eCMR builds on the Context Maintenance and Retrieval Model (CMR, Polyn et al., 2009), but extends this model by implementing encoding priority as increased context-to-item associations in a two-layer neural network. After encoding of a mixed list, when the memory test relies on the context to cue retrieval (i.e. in a free-recall test, when no other cues are available), these stronger associations promote the recall of items that were prioritised during encoding. Once a high-reward item is retrieved, it will reactivate the high-reward context of its encoding, thereby promoting the recall of items that share the same context (other high-reward items). A second aim of this study was to test specific behavioural predictions of eCMR. Our second and third hypotheses are that high-reward items are retrieved first in mixed lists, and that recall would be clustered according to the reward condition. These hypotheses were not pre-registered, but are derived directly from the eCMR model (Talmi, Lohnas, et al., 2019).

A central assumption of the way eCMR simulates encoding of high-reward items is that they are more tightly-bound to their encoding context, a mechanism considered to be a computational implementation of context-independent increases in the attention allocated to prioritised items, and to operate during encoding regardless of list composition. We recorded EEG during encoding with the aim to test this assumption and more broadly to investigate the neural mechanisms underlying reward-enhanced memory. We investigated EEG measures that have been related to attention: steady-state visual evoked potentials (ssVEPs) and event-related potentials (ERPs), including P3, LPP (Late Positive Potential). Our fourth, pre-registered hypothesis is that the amplitude of ssVEP and ERP effects will be larger for high than low reward items regardless of list composition, namely, observed in both mixed and pure list conditions, reflecting list context-independent increased attention to high reward items.

The ssVEP technique has been used extensively to study visual attention (for reviews see (Vialatte et al., 2010; Wieser et al., 2016). The technique involves flickering stimuli at a given frequency; increased visual attention is associated with increased ssVEP power in the flicker frequency over occipital electrodes. ssVEP amplitude is increased by reward (e.g. Grahek et al., 2021) and predicts subsequent memory (Martens et al., 2012). ssVEPs and ERPs have traditionally been used as indices of attention in separate literatures, but it is possible to obtain both measures in the same study (e.g. Hajcak et al., 2013; Jiang et al., 2018; Thigpen et al., 2018). The P3 component and the late posterior positivity (LPP) have both been related to increased attention during encoding. The P3 component is related to allocation of attention and reward processing (Polich, 2007; Sato et al., 2005) and has recently been shown to be higher for high-vs. low-reward items during encoding (Elliott et al., 2020). The P3 is a positive peak 250-500ms post stimulus presentation which is maximal over midline parietal electrodes (Polich, 2007). The LPP has traditionally been associated with allocation of attention to emotional compared to neutral stimuli (for review see Hajcak et al., 2010; Schupp et al., 2000, 2006), but it may also more broadly reflect stimulus significance and may thus be sensitive to reward (Hajcak & Foti, 2020). The LPP is a positive slow wave, which is maximal over centroparietal electrodes and onsets around 200-400ms after stimulus presentation and typically lasts as long as the emotional stimulus is presented (Hajcak & Foti, 2020; Schupp et al., 2006). Two studies reported that the emotional modulation of the LPP is insensitive to list composition, and one found that it was related to subsequent memory (Barnacle et al., 2018; Zarubin et al., 2020). In addition to the ERP effects related to attention we also investigated the frontal slow wave (FSW) ERP effect, which has been related to elaborative rehearsal strategies (Fabiani et al., 1990; Mangels et al., 2001). The FSW typically onsets late, around 1000ms post stimulus presentation and has a frontal maximum (Elliott et al., 2020). Crucially, because ssVEPs and ERPs, including the FSW, are time-locked to stimulus presentation, their modulation by reward suggest that reward influences attention to and rehearsal of valued stimuli during stimulus presentation. Any one of these effects could implement reward-dependent tightening of the association between the context and item layer in eCMR.

The eCMR account of list-composition effects can be seen as an instantiation of the retrieval distinctiveness account (Geraci et al. 2013; McDaniel et al. 2005). Retrieval distinctiveness accounts of list composition effects attribute the effect to prioritization of reward items at encoding, which endows them with properties that render them more distinctive during retrieval, and therefore better retrieved than less-distinctive items during the recall of mixed, but not pure lists. In eCMR, increased context-to-item binding provides a specific mechanism that renders particular items (here, those associated with high reward) ‘stronger’ than ‘standard’ items (here, those associated with low reward). eCMR can also be contrasted with two other accounts of list-composition effects. The item-order account (McDaniel & Bugg, 2008) suggests that reward captures encoding resources otherwise dedicated to encoding temporal order, and therefore predicts that temporal order will be less well preserved in the recall of mixed lists and pure, high-reward lists compared to pure, low-reward lists. Although eCMR assumes that reward items are bound more strongly to their context, the portion of context that they are bound more strongly to is the partition that represents reward rather than time. In technical terms, in eCMR high-reward items are more strongly bound to the “source context” (Polyn et al., 2009), rather than the temporal context (Talmi, Lohnas & Daw, 2019), and consequently, it does not predict that the composition of the list or the reward value will influence temporal organisation. Therefore, if temporal organisation scores change as a function of list type and reward, and especially if the direction of change follow that predicted by the item-order account, this result will support the item-order account and challenge eCMR. We have carried out a similar analysis for the emotion-dependent list-composition effect, which favoured the eCMR account, and therefore, although we explore this again here, we have not registered this hypothesis. Finally, the attentional borrowing account attributes list-composition effects to selective rehearsal of high-reward items. These ‘borrows’ attention from other items in mixed lists but in pure lists they cannot do so (Watkins, LeCompte & Kim, 2000; Slamecka & Katsaiti, 1987). Attention borrowing can take place immediately upon item presentation and shortly after (akin to massed rehearsal), or during the presentation of other items (akin to spaced rehearsal). While the former is similar to the extra attention capture that eCMR already simulates, eCMR cannot account for selective spaced rehearsal of high-reward items. Under natural encoding conditions it is tricky to pinpoint exactly when people rehearse. However, if selective spaced rehearsal is the only reason for list-composition effects in free recall, there should be no effect of reward on EEG marker of attention in pure lists. This prediction contrasts directly with the eCMR account of the list composition effect, and corresponds to our fourth pre-registered hypothesis, as described above.

The present study was the first to investigate the neural mechanisms underlying the context dependence of reward-enhanced memory. By combining multiple EEG measures of attention and selective rehearsal the study tests the relative contribution of automatic reward signals and meta-cognitive encoding strategies in reward enhanced memory and specifically tests the assumption of increased attention during encoding of the eCMR model.

## Methods

### Participants

The sample size was determined a priori based on effect sizes in Hajcak et al. (2013). Power analysis using the WebPower package (Zhan & Mai, 2018) in R (v4.0.5; R Core Team 2021) indicated that a sample size of 43 is needed to replicate their ssVEP main effect of emotion (*η*^2^*_p_* = .203) with .90 power at alpha = .05. The data was collected using a pre-registered stopping rule stating that data collection would continue until the targeted sample size was reached and that excluded participants based on pre-registered exclusion criteria would be replaced. Fifty-one participants gave informed consent before participating in the experiment. The participants were compensated with £25 for their participation plus the performance-based reward earned in the experiment. The reward was calculated for two randomly selected lists plus the practice list (the maximum total reward was £16.8). Eight participants were excluded based on pre-registered criteria: three based on having less than 16 trials in one of the four conditions after EEG pre-processing, three due to not having an ssVEP at 14.17Hz when averaging over all four conditions, one participant due to a technical error and another participant due to performing below the 75% threshold in the distractor task. The final sample consisted of 43 participants (31 female, 11 male, 1 non-binary; *M*_age_ = 24.4, *SD*_age_ = 4.6). The experiment was approved by the Cambridge psychology research ethics committee.

### Materials

The stimuli used in the study were 144 images (585x410 pixels) obtained from a standardized set of 260 stimuli (Snodgrass & Vanderwart, 1980). The assignment of picture to condition was randomised. There were three different types of lists: pure high reward (two lists), pure low reward (two lists) and mixed lists (four lists).

Two coloured frames surrounding the image were used to indicate the reward condition. We used a photometer to control their luminance. The selected colours were the following in RGB space, brown 101, 55, 0 with 100% opacity; blue 3,67, 223 with 86% opacity. The assignment of colour to reward condition was counterbalanced across participants. The pictures were transformed to grayscale and presented on a 19 inches CRT monitor with a refresh rate of 85Hz.

### Experimental Design

We used a 2 (Reward: High vs Low) by 2 (List composition: mixed list vs pure list) design. There were nine blocks. The first block was a practice block. Four of the blocks had mixed lists with both high and low reward items whereas the other four lists were pure and either only had high reward items (two lists) or only low reward items (two lists). The assignment of list type (mixed, pure/low reward, pure/high reward) to block was randomised.

### Procedure

Each block of the experiment started with an encoding phase. In this phase, the participant was shown 16 pictures sequentially in the centre of the screen. Each trial started with a fixation cross with a randomised jittered duration between 3.5, 4 or 4.5 seconds. Next a picture was flickered at 14.17Hz in the centre of the screen for 2047 ms. The picture had a coloured frame (blue or brown) which indicated what reward condition it was associated with. The participants were instructed that they would receive £0.5 for remembering pictures from the high-reward condition (e.g. pictures with blue frames) and £0.05 for remembering pictures from the low-reward condition (e.g. pictures with brown frames).

A one-minute distractor task followed the encoding phase. A math problem (e.g. 5 + 9-3 = 11) was shown on the screen and the participants were asked to indicate via button presses if the answer was true or false. We told the participants that their responses needed to be 75% correct in this distractor task to get the monetary reward for the memory task. All included participants reached this criterion.

A free recall test followed the distractor task. The participants had three minutes to write down descriptions of the pictures they could recall at any order, using a keyboard. They were encouraged to "use multiple words or phrases to describe one picture, e.g. ’spotty dog’ and ’cat lying down’ as opposed to ’animal’; ’lettuce’ and ’broccoli’ as opposed to vegetable" and to "be as specific as possible so we know which picture you are referring to. The participants were encouraged to continue to try to recall until the end of the response time window. The last response was on average entered after 111 seconds (*SD* = 23 seconds), indicating that the participants had enough time respond.

### Behavioural data analysis

Accuracy in the final memory test was coded manually by the experimenter (RH) who compared participants’ responses to the pictures shown in each list while being blind to experimental conditions. Fifty-one percent of the data (22 out of the included 43 subjects) were coded by a second coder and the percentage of correspondence between the two coders were 98.2% for this proportion of the data. The pre-registered analysis of memory performance utilised a generalised mixed effect model using the lme4 package (Bates et al., 2015) in R. Memory performance was used as a dependent variable and there were two fixed effects of Reward (high vs low) and List composition (mixed lists vs pure lists) and a random intercept plus random slope effect of participant. The model formula was:

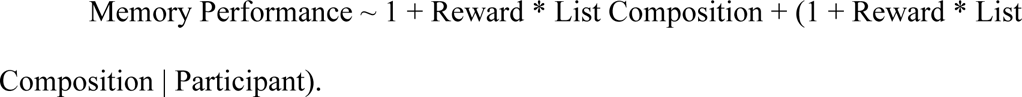

The statistical significance of the fixed effects and their interaction was assessed using an omnibus χ^2^ Wald test as implemented in the car package (Fox & Weisberg, 2019). We followed recommendations in (Brown, 2021) to handle convergence issues (i.e. we changed the optimiser and increased the number of iterations for the memory performance GLMM).

According to hypothesis 2, high reward items are more likely to be recalled in the first output position than low reward items. An analysis of this effect must control for the expected increase in recall of high reward items. Therefore, for each mixed list, we permuted the output positions of recalled items 10000 times, to estimate the chance level in our data when accounting for the increase in recall for high reward items. We then compared the chances of recalling a high-reward item in the first output position in the observed and the permuted data with a one sample test (for a similar approach see Healey et al., 2019). A Wilcoxon signed rank test was used due to that that the Shapiro-Wilk test of normality was significant (*W* = .838, *p* < .001), suggesting deviation from normality.

Next, we tested whether the participants organised recall based on reward condition (hypothesis 3). Again, we controlled for clustering that could occur by chance due to higher memory performance in the high reward than in the low reward condition by permuting the output order 10000 times. We calculated the reward-based organisation scores for each participant in the observed and the permuted data using the EMBAM toolbox (https://github.com/vucml/EMBAM) in MATLAB. Observed reward-based organisation scores were compared to the mean score in the permuted data using a one sample test. A Wilcoxon signed rank test was used due to that that the Shapiro-Wilk test of normality was significant (*W* = .935, *p* = .018), suggesting deviation from normality. Finally, to contrast eCMR and the item-order account, we used the EMBAM toolbox to compute temporal organisation scores. We then conducted a one-way repeated measures ANOVA with temporal organisation score as dependent variable List Type as independent variable (mixed list, pure list high reward, pure list low reward). Given that eCMR predicts no difference in temporal organisation scores between list types we complimented the frequentist ANOVA with a Bayesian one-way repeated measures ANOVA conducted in JASP (JASP Team 2022, version 0.16.3).

### EEG recording and pre-processing

EEG data was acquired using a 128-channel high-density hydrocel Geodesic sensor net, sampled at 500Hz using a Netamps 300 amplifier. The data was recorded during the covid-19 pandemic and the participants were wearing face masks throughout the experiment which necessitated excluding the 15 electrodes covering the face. The EEG data was pre-processed offline using the EEGLAB toolbox in MATLAB with a standard pipeline (see Hellerstedt et al. 2021 for a study using a similar pipeline) and the steps and settings were pre-registered. The data was recorded with Cz as reference and was re-referenced offline to the average of the left and the right mastoid electrodes. Next, the data was high-passed filtered with an FIR filter with a cut off at 0.1Hz and segmented into epochs starting 2000ms prior to stimulus onset and ending 3000ms after stimulus onset. Channels and epochs with artefacts were detected and rejected using customised threshold functions from the FASTER plugin (Nolan et al. 2010). The criterium for rejection of channels were values outside ±3 for participant-level *z*-transformed values. We used the standard settings in FASTER meaning that the values used for epoch rejection were the amplitude range, variance and deviation from channels mean and the values used for rejecting channels were each channel’s mean correlation with other channels, variance and hurst exponent, a measure of a signal’s long-range dependence. Next the data was submitted to independent component analysis and independent components reflecting ocular artefacts were identified by visual inspection of scalp topographies, time course and activation spectra and were discarded from the data by back-projecting all but these components to data space. Rejected channels were interpolated and a low-pass FIR filter was applied with a cut-off at 40Hz. Channels containing artefacts within single epochs were detected using customised functions from the FASTER plugin. The criterion for rejecting and interpolating a channel was that *z*-transformed values exceeded ±3 within a single epoch. We used the standard settings in FASTER meaning that the values compared against the threshold were variance, median gradient, amplitude range and channel deviation (Nolan et al., 2010). Epochs with amplitudes exceeding ±100 microvolts were detected and rejected to clean out any remaining artefacts. Finally, the epochs were sorted into the conditions. Mean trial numbers ranged between 28.9 and 29.3, out of 32, for the four conditions and all participants contributed with at least 16 trials per condition.

### Focal ssVEP analysis

The procedure for the focal ssVEP analysis was pre-registered. The EEG data was baseline-corrected using a pre-stimulus interval of 500ms. Next, the power spectrum was computed for the time-window between 1000-2000ms post stimulus presentation using Fast Fourier Transform with a Hanning taper as implemented in the FieldTrip toolbox (Oostenveld et al., 2011) in MATLAB (ver. 2021a). Amplitude values were extracted between 14Hz and 15Hz, the stimulation frequency, from electrode Oz. The selection of time window and electrode of interest was based on where and when the ssVEP was maximal in time-frequency data when averaging over all four conditions as described in the pre-registration. Extracted ssVEP amplitude data were analysed using linear mixed effects models using the lme4 package (Bates et al., 2015) in R. The extracted ssVEP amplitudes were used as the dependent variable. We started out from a model including a fixed effect of Reward (High/Low) a fixed effect of List composition (Mixed list/Pure list) and random intercept + random slope effects of participant, but needed to reduce the complexity of the random effect structure to only include random intercepts due to convergence problems. The formula for the final model was:

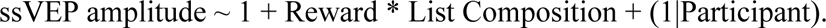

### Global ssVEP analysis

Analysing EEG data from a selected time-window and region may overlook effects occurring at other scalp locations and time points. We therefore complemented the focal analysis with a pre-registered global data driven analysis of all time points of the trial, 0-2000ms after stimulus presentation, and all 113 analysed electrodes and controlled for multiple comparisons with nonparametric cluster-based permutation tests with the FieldTrip toolbox (Maris & Oostenveld, 2007).

First, single trial EEG-data was submitted to time-frequency decomposition using Fast Fourier Transform with Hanning taper. The decomposition was performed between 4 and 30Hz in frequency steps of 1Hz, time steps of 50ms and with a time-window length of five cycles. Oscillatory power in each frequency band was normalised to dB scale using a pre-stimulus baseline period between –700 to –200ms to avoid that the estimation of baseline data points were affected by data in the post-stimulus time window in the stimulation frequency (as in Allen et al., 2020; Hellerstedt et al., 2021).

Next, nonparametric cluster-based permutation tests were performed for data in the stimulation frequency between 14-15Hz. In the first step of this analysis t-tests are performed for every timepoint at every electrode. Significant spatially or temporally neighbouring samples (including at least two electrodes) at alpha = .05 (uncorrected) were used to form clusters and their t-values were summed to a cluster-level *t*-value. The observed cluster-level *t*-value was then compared to the maximal cluster-level *t*-value obtained in 5000 permutations. The Monte-Carlo *p*-value was calculated as the proportion of the permutated cluster-level *t*-values being larger than the observed cluster-level *t*-value. The exact spatiotemporal distribution of the clusters should be interpreted with caution given that the correction for multiple comparisons is applied at the cluster-level, but not at the individual sample level (Sassenhagen & Draschkow, 2019). We tested the main effect of Reward, the main effect of List composition and the Reward x List composition interaction.

### Global ERP analysis

Two notch filters were applied to filter out the ssVEPs from the EEG data prior to ERP analysis. The first filter had a lower cut-off at 14Hz and a higher threshold at 15Hz to filter out the stimulation frequency and a second filter had a lower cut-off at 28Hz and a higher cut-off at 29Hz to filter out the resonance frequency. The ERP data was baseline corrected using a pre-stimulus interval between –200ms and 0ms. As for the global ssVEP analysis, the global ERP analysis used nonparametric cluster-based permutation tests of all timepoints in the trial from 0-2000ms and included all 113 electrodes and the analysis was pre-registered. The settings were identical to the global ssVEP analysis.

## Results

### Behavioural Results

Figure 1A depicts the proportion of items recalled in each of the four list-composition by reward conditions. We begin by reporting results that correspond to the pre-registered hypothesis 1 and the pre-registered analysis. There was a reliable main effect of Reward, (χ^2^(1) = 20.56, *p* < .001) while the main effect of List composition was not reliable (χ^2^(1) = 1.57, *p* = .21). Importantly, the Wald test also indicated that the Reward x List composition interaction was reliable (χ^2^(1) = 17.24, *p* < .001). Consistent with our pre-registered prediction, follow-up pairwise contrasts showed that memory performance was higher for high reward than low reward items in mixed lists (z-ratio = –5.200, *p* < .001), but not in pure lists (z-ratio = –1.716, *p* = .086), indicating that reward enhanced memory depended on list context. Parameter estimates of the model are presented in Table 1.

**Figure 1.**
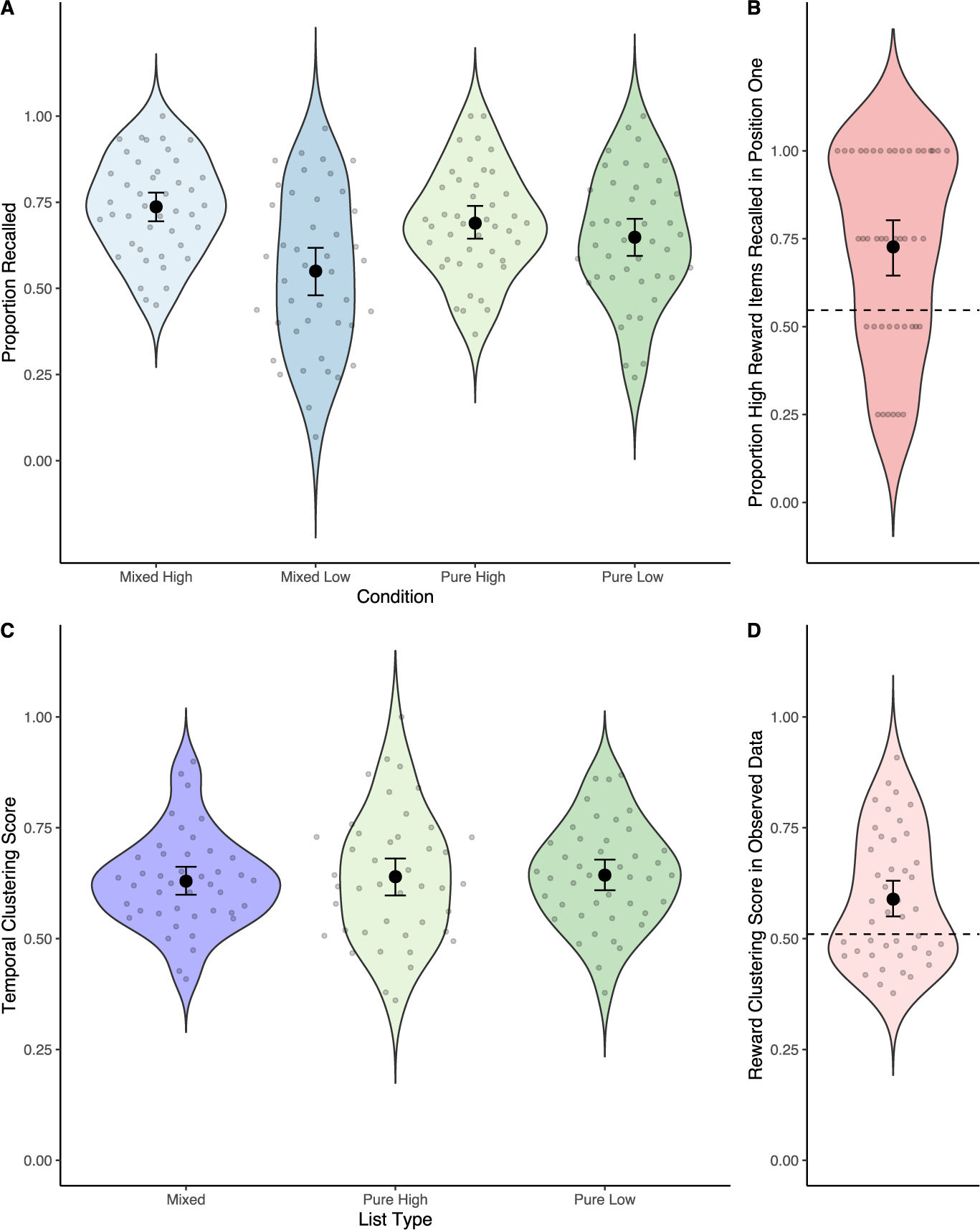
Behavioural results. A) Proportion recalled items in the final memory test for the four conditions.Scatter points indicate means for each participant. The larger black dots displays the group means and the errorbars reflect the 95% confidence interval of the group means. B) Proportion of high reward items recalled first in the mixed lists in the observed data. The dotted line shows the mean proportion of high reward items recalled first in the permuted data. C) Temporal clustering scores for the three list types. D) Clustering score in the observed data. The mean clustering score in the permuted data is displayed as a dotted line.

**Table 1.** Model parameters for memory performance GLMM and the targeted ssVEP LMM analysis.

| | Estimate | SE | $z/t^*$ |
| --- | --- | --- | --- |
| Memory Performance GLMM |  |  |  |
| Intercept | .75 | .11 | 6.60 |
| Reward | -.27 | .06 | -4.53 |
| List composition Pure | -.05 | .04 | -1.25 |
| Reward:List Composition Pure | -.17 | .04 | -4.15 |
| ssVEP LMM |  |  |  |
| Intercept | 3.52 | .37 | 9.32 |
| Reward | .04 | .06 | .77 |
| List composition Pure | -.07 | .06 | -1.15 |
| Reward:List Composition Pure | -.02 | .06 | -.28 |

\*Z for GLMM and t for LMM

Next, we examined two additional hypotheses which were not pre-registered, but derived from the eCMR model. Participants were more likely to recall high reward items first in the mixed lists. To test whether this reflected a bias to recall high-reward items first, or was a consequence of the overall increased recall of high-reward items, we compared observed and permuted data. A one sample Wilcoxon signed rank test indicated that the likelihood of retrieving a high reward item in position one was reliably higher in the observed data (*M* = .727) than in the permuted data (*M* = .547; *V* = 760, *p* < .001; see Figure 1B). These results support hypothesis 2, suggesting that participants were more likely to retrieve a high reward item first, even considering the general effect of reward on recall. To test hypothesis 3 we analysed the likelihood that output order was organised based on reward condition. This analysis queries whether participants were more likely to retrieve a high reward item in a given trial if they retrieved a high reward item in the preceding trial. Confirming hypothesis 3, a Wilcoxon signed rank test showed that reward clustering was greater than the estimated chance level (*M* observed = .589, *M* permuted = .510; *V* = 702, *p* =.005; see Figure 1D).

Finally, we assessed the effect of reward and list composition on temporal organisation scores (Figure 1C). This analysis examines whether participants were more likely to retrieve items that they have studied consecutively during encoding. A one-way repeated measures ANOVA indicated that there was no effect of list type (mixed list vs pure list high reward vs pure list low reward) on temporal organisation scores (*F*_2,84_ = .258, *p* =.773, η_p_^2^ = .006) and a Bayesian one-way ANOVA showed strong evidence for the null hypothesis over the alternative (BF01 = 10.792). These results are in line with predictions from eCMR and difficult to reconcile with the item-order account.

### ssVEP manipulation check

The frequency spectrum of the EEG was extracted for the whole trial (0-2000ms) averaged over all conditions to check that the flicker manipulation had induced an ssVEP at 14.17Hz using a fast fourier transform (FFT) with a Hanning taper. These data confirm that the flicker manipulation successfully induced increased spectral power at 14Hz and in the resonance frequency at 28Hz (Figures 2A, 2B and 2C). We next performed a pre-registered time frequency analysis to detect what time window to use for the focal ssVEP analysis. In line with previous research, this analysis showed that the ssVEP was maximal over occipital electrodes and between 1000-2000ms post stimulus presentation (Figures 2C and 2D).

**Figure 2.**
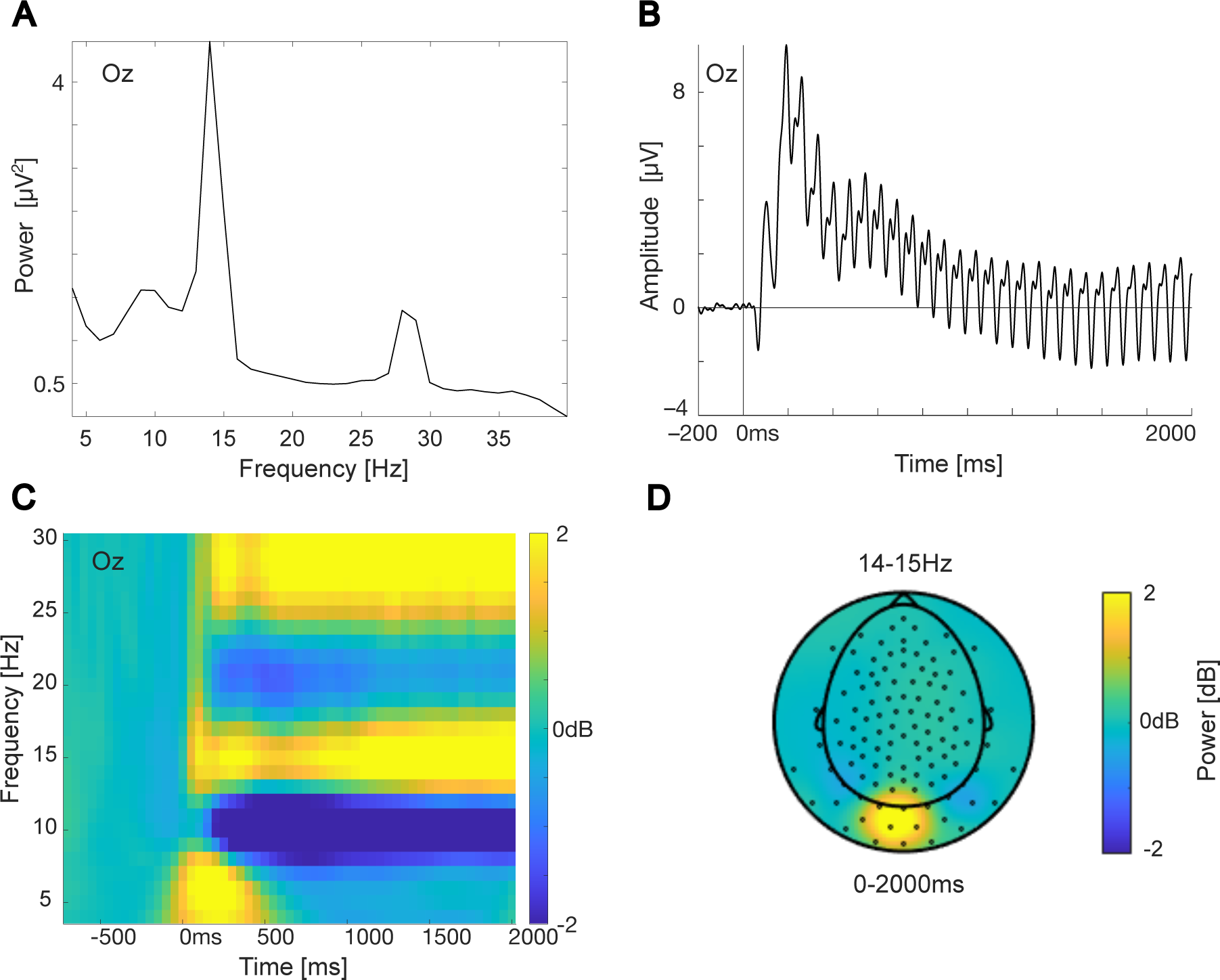
ssVEP Manipulation Check. A) Power spectrum of all conditions combined from electrode Oz showing increased power in the stimulated frequency at 14Hz and at the resonance frequency at 28Hz. B) ERP waveform averaged over all trials from Oz showing the ssVEP at 14Hz. Note that the ssVEP was reducedwith notch filters prior to the statistical analysis of ERPs. C) Time frequency plot averaged over all four conditions from electrode Oz showing that the increase in power at 14Hz was maximal around 1000-2000ms. D) Topographic map over time frequency data between 14-15Hz for the whole epoch averaged over all conditions and averaged over the whole stimulus time window (0-2000ms).

### Focal ssVEP analysis

Figure 3 depicts the pre-registered focal ssVEP analysis. Contrary to predictions, there was no main effect of Reward, χ^2^(1) = .60, *p* = .440, or list composition χ^2^(1) = 1.32, *p* = .250, and the interaction between Reward and List composition, χ^2^(1) = .08, *p* = .782 was also not reliable, indicating that there were no effects of Reward or List composition on ssVEP power. We had pre-registered a focal ssVEP subsequent memory analysis, to investigate if any observed effects were related to memory, but we did not conduct this analysis due to low trial numbers when dividing the EEG trials for the four conditions based on performance.

**Figure 3.**
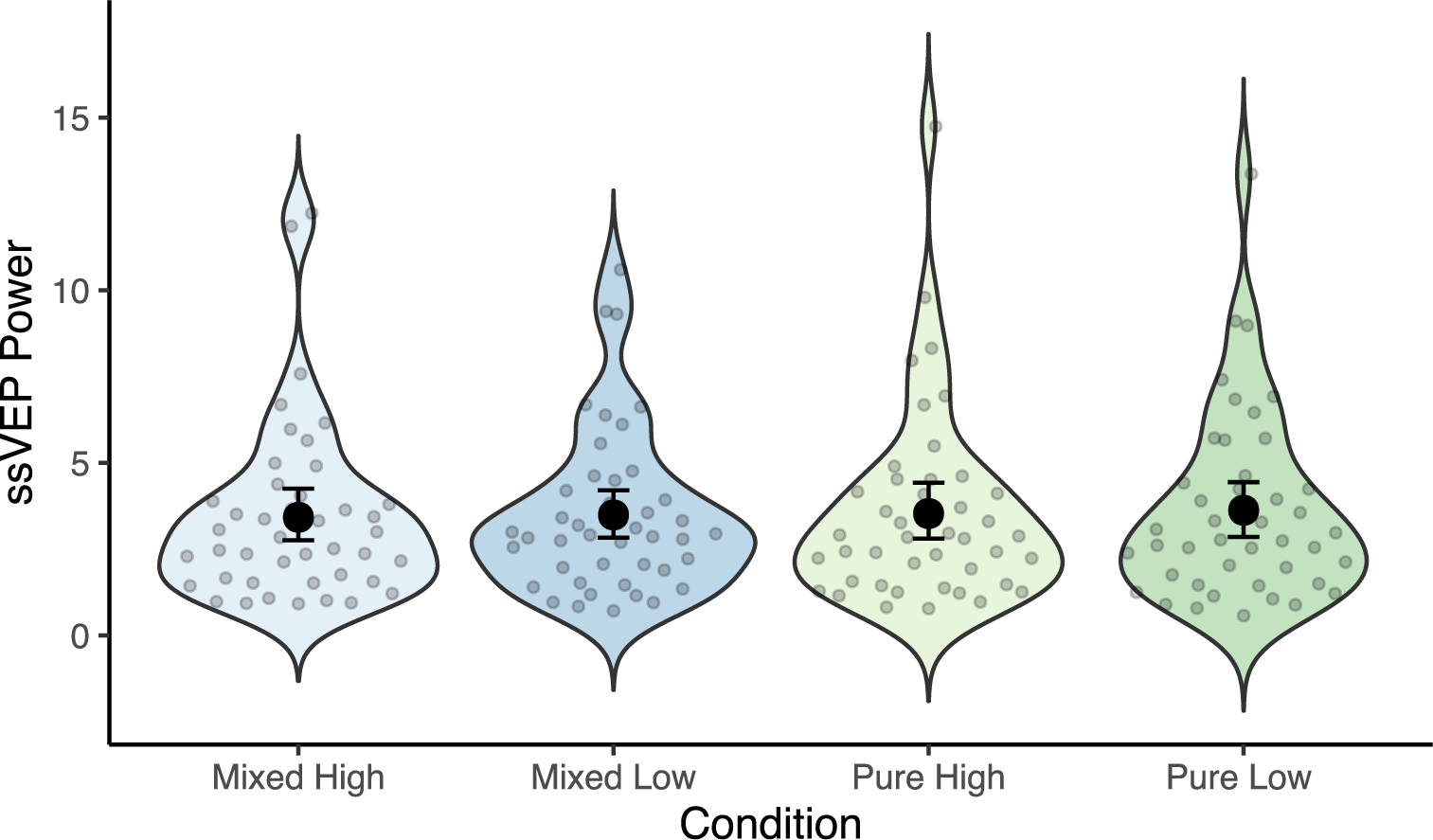
Focal ssVEP analysis. The scatter dots depict individual ssVEP power between 1000-2000ms post stimulus presentation from electrode Oz for the four conditions. The larger black dots reflects group means and the error bars shows the 95% confidence interval of the group mean.

### Global ssVEP analysis

Figure 4 depicts the pre-registered exploratory global analysis. There was a reliable negative cluster in the analysis in the main effect of Reward, showing larger ssVEP power for low reward than for high reward (*p* = .003). The cluster was reliable between approximately 750-1650ms and was maximal over occipital and parietal electrode sites. There was no reliable main effect of list composition (all *p*s > .724) and no reliable Reward x List composition interaction (all *p*s > .421). Although we have predicted to find a main effect of reward, the direction of this effect was surprising and contradicted our pre-registered hypothesis, as discussed in the Discussion section below.

**Figure 4.**
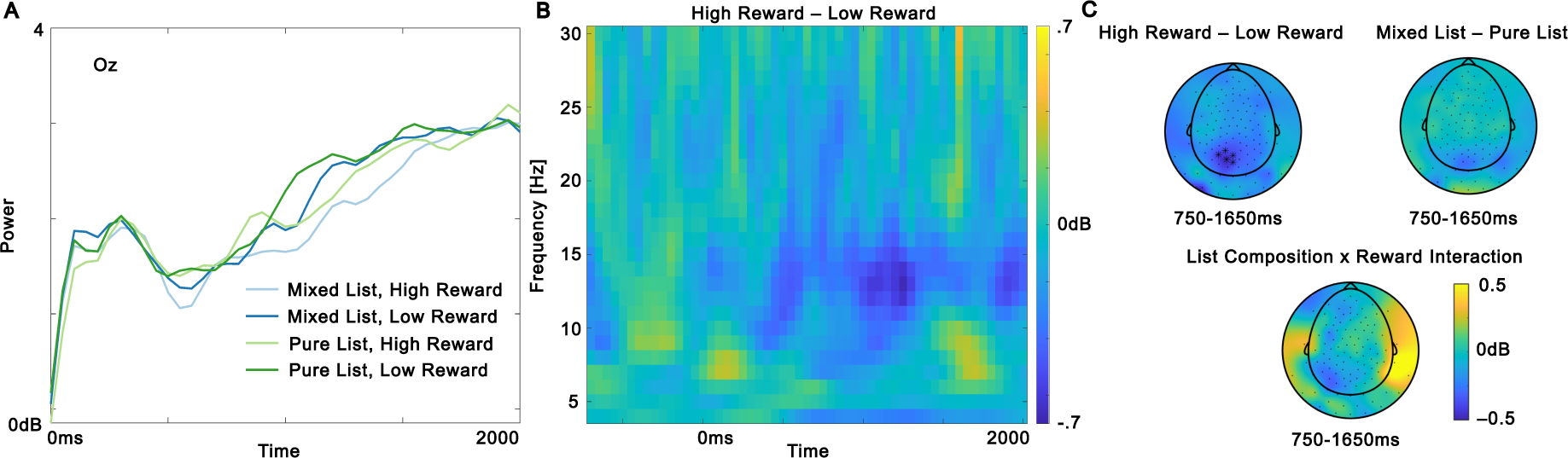
Global ssVEP analysis. A) Time frequency data from electrode Oz between 14 and 15Hz for the four conditions. B) Time-frequency plot with the averaged power difference between high reward and low reward from the five electrodes in the significant cluster in the global ssVEP analysis. C) Topographic maps of power differences between 14-15Hz for the time window where there was a reliable negative cluster for the main effect of reward. Electrodes belonging to a significant cluster which are significant on average in the displayed time window are highlighted with black stars.

As can be seen in Figure 2C, the experimental task was overall associated with a reduction in alpha power around 10Hz and in beta power around 20Hz when averaging over all conditions. We conducted control analyses to investigate if the unexpected reduction in ssVEP power for high reward items compared to low reward items was driven by a global reduction in alpha/beta power. If the effect was driven by global alpha/beta suppression, we would expect to see a similar effect outside the stimulation frequency. To investigate this possibility, we performed cluster permutation tests including all electrodes and all time points for the alpha band (8-12Hz, *p* = .169) and the beta band (13-30Hz, *p* = .152), but found no significant clusters in these frequency bands. We also tried to analyse narrow frequency bands just below (11-13Hz, *p* = .05, which was not significant given the statistical threshold of *p*=.025 for the two tailed test) and just above (15-17Hz, *p* = .267) the stimulation frequency. The results suggest that the effect was most robust in the ssVEP frequency and therefore, was unlikely to be driven by alpha/beta suppression. Figure 4B illustrates that the decrease in power for high reward compared to low reward was maximal around the ssVEP frequency.

### Global ERP analysis

ERP results from the pre-registered global analysis are depicted in Figure 5. There was a reliable positive cluster for the main effect of Reward (*p* = .009) between 1050-2000ms with a right anterior/central distribution, indicating greater ERP amplitude for high reward than low reward. There was no main effect of List composition (all *p*s > .130). Mirroring the behavioural results, there was a reliable Reward x List composition interaction (*p* = .014) which was reliable around 860-1320ms and had a posterior and central distribution. Follow-up analysis showed that there was a reliable positive cluster around 590-1450ms within the mixed lists, indicating more positive amplitude for high reward items than low reward items with a widespread topography (*p* = .001). The effect has a posterior topography around 600-1000ms fitting with an LPP effect (e.g. Barnacle et al., 2018; Hajcak et al., 2013; Zarubin et al., 2020) while it has a more anterior topography after 1000ms fitting with the spatiotemporal characteristics of an FSW effect (e.g. Elliott et al., 2020). The effect is more long-lived than a typical P3 effect which is a peak rather than a slow wave, although there are ongoing discussions regarding whether the P3 and the LPP are neural reflections of the same processes and whether the duration of the effect may be determined by tasks characteristics (Hajcak & Foti, 2020). There were no reliable clusters when comparing amplitude for high vs low reward within pure lists (all *p*s > .837).

**Figure 5.**
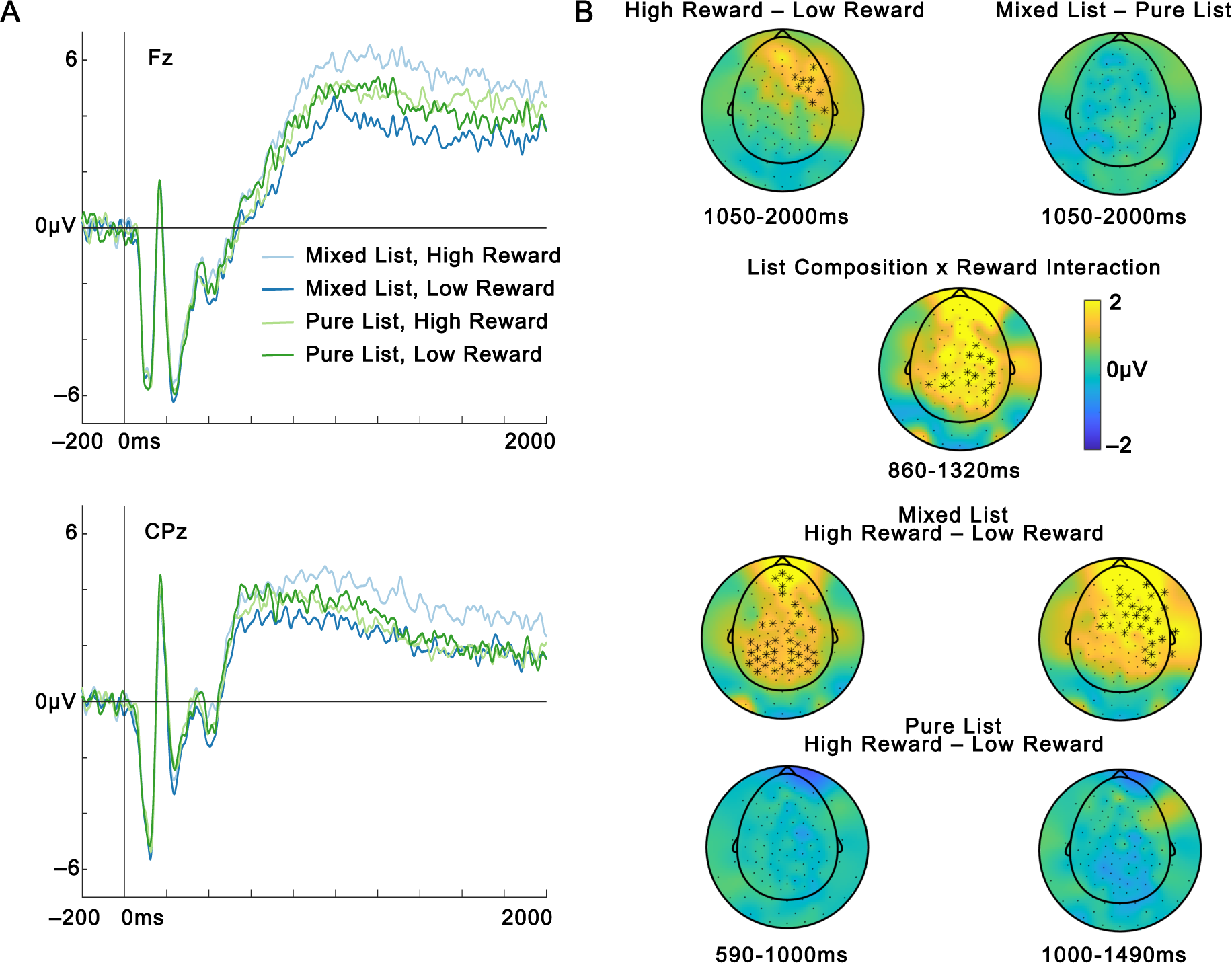
Global ERP analysis. A) ERPs from electrodes from two electrodes where the ERP effects of interest typically are maximal, a frontal electrode (Fz) for the FSW and a centroparietal electrode (CPz) for the LPP. B) Topographic maps of amplitude differences from the time windows of the reliable clusters for the different contrasts. The cluster for the follow up of the List Composition x Reward interaction, the bottom four plots, has been divided into an early (590-1000ms) and a late (1000-1490ms) part to show the change in topography over time. Electrodes from reliable clusters are highlighted with black stars if they are reliable on average in the plotted time window.

Scalp current source densities (CSDs) were computed using the spherical spline method (Perrin, Pernier, Bertrand & Echallier, 1989) as implemented in FieldTrip to further investigate if there was support for an anterior and an posterior source for the reward ERP effect within mixed lists. As depicted in Figure 6 the CSDs provided support for both a posterior and an anterior source in the 590-1000ms time window (left plot). This is the time window when the LPP typically is maximal, so this is in line with the interpretation that a posterior LPP ERP effect and an anterior FSW ERP effect are present in this time window. The FSW is typically maximal after 1000ms and the LPP is typically reduced then which also matches the CSD pattern where the posterior source gets weaker while the anterior source continues to be strong in the 1000-1490ms time window.

**Figure 6.**
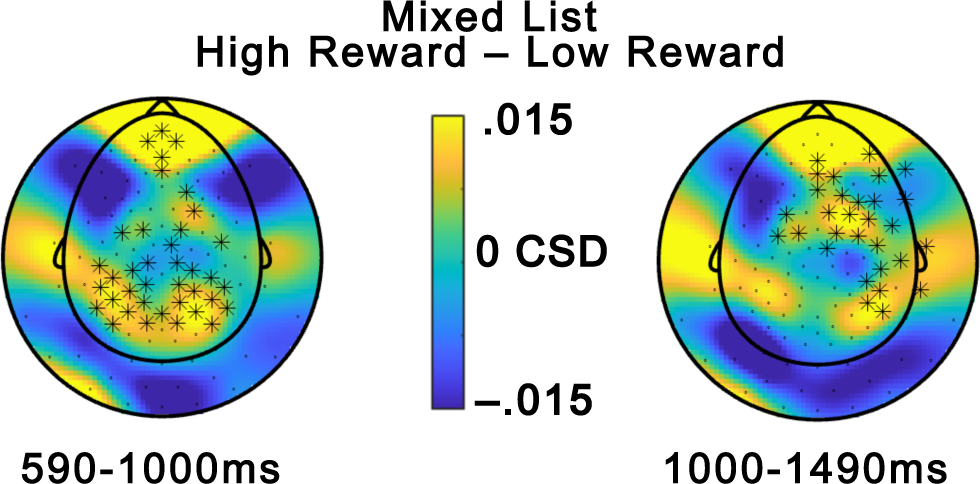
Current source density plots from an early and a late time window from mixed lists where there were significant differences between high- and low-reward ERP amplitudes. Electrodes from reliable clusters are highlighted with black stars if they were reliable on average in the plotted time window.

## Discussion

The present study investigated whether reward-enhanced memory is context-dependent, and the neurocognitive mechanisms underlying reward-enhanced memory. We also tested predictions of the eCMR computational model of memory. All four behavioural predictions of eCMR were supported by the data. By contrast, the EEG data challenged our proposed neural implementation of a key eCMR mechanism. The ERP results showed for the first time that the modulation of LPP and ssVEP by reward is context dependent, suggesting that they reflect top-down controlled strategic processes that are applied when high and low reward items compete for cognitive resources. In contrast to the ERPs, the ssVEPs differentiated between high and low reward in a context-independent manner, potentially reflecting increased focus on internal semantic representations relative to external visual perceptual information when encoding high reward items compared to low reward items. Taken together, the pattern of the EEG results show that LPP and ssVEPs are functionally dissociated, suggesting that they measure different aspects of attention.

Reward-enhanced memory was dependent on list context, meaning that recall of items that predicted reward was increased in mixed lists, but not in pure lists, conceptually replicating two recent studies from our group (Hellerstedt & Talmi, 2022; Talmi et al., 2021) despite a number of methodological differences (e.g. nature of the stimuli). The finding that reward did not enhance recall in pure lists places an important constraint on any explanation of reward-enhanced memory as solely a function of automatic encoding processes. It aligns better with other explanations: either the eCMR model and other retrieval-distinctiveness accounts, where encoding-retrieval interactions play a key role, or the suggestion that automatic processes operate together with strategic processes to enhance memory for valued information (Hennessee et al., 2019; Knowlton & Castel, 2022).

Retrieval distinctiveness accounts explain the list-composition effect as a consequence of prioritised encoding, which is expressed differently during the retrieval of mixed and pure lists (Gerarci et al. 2013; McDaniel et al 2005). eCMR instantiates this account with a computational mechanism that makes predictions about recall dynamics. Findings from the mixed list condition agreed with the recall dynamics that this computational model predicted. In agreement with the prediction that the high reward items are more tightly bound to the encoding context and therefore will be more likely to be recalled first (Talmi, Lohnas, et al., 2019; Talmi et al., 2021), participants recalled high-reward items earlier than low-reward items. While in previous studies of recall of mixed lists of ‘strong’ and ‘weak’ items, including ours, the propensity to recall ‘stronger’ items early was confounded with the increased average recall of such items, here we controlled for this confound yet still observed that high-reward items were recalled earlier than low-reward items. Additionally, we found that recall of mixed lists was organised based on reward condition. This finding confirms a second prediction of eCMR, that recalling a high reward item will change the context of the recall test to overlap more with the high-reward encoding context, and therefore, promote recalling further high-reward over low-reward items. Finally, we obtained evidence that neither reward, nor list composition, alters the temporal organisation of recall. This finding supports eCMR, where high-reward items are bound more tightly to their source context (the context partition that codes reward) but not their temporal context. By contrast, this finding challenges the item-order account of list composition effects (McDaniel & Bugg 2008; Nairne et al, 1991). While recall dynamics qualitatively align with predictions of eCMR, this is not conclusive evidence that this model is correct or detract from a potential role of meta-cognitive strategic encoding processes in reward-enhanced memory. For example, a model that formalises spaced rehearsal processes, e.g. selective rehearsal of high reward items in working memory across the entire presentation of mixed lists, may offer a better fit. EEG data offer additional insight into the contributions of automatic reward processes, meta cognitive processes, and the specific mechanisms of eCMR in reward-enhanced memory.

ssVEPs offer an unobtrusive measure of the amount of visual attention participants allocate to items during encoding. eCMR embodies tighter associations between the context and highly-attended items regardless of the local context, and we have therefore pre-registered a hypothesis that if ssVEPs indexes this effect, ssVEP power will be increased in the high-reward condition. In contrast, the attentional borrowing account postulates that high-reward items borrow attentional resources or rehearsal time during encoding from low reward items when they compete for attention in mixed lists, but not in pure lists (Watkins, LeCompte & Kim, 2000). The ssVEP results agreed with the prediction from eCMR in that they showed a context-independent effect of reward. However, surprisingly, we observed *a reduction* in ssVEP power for high-reward items, suggesting decreased visual processing of these stimuli. One plausible interpretation of this result is that participants focused more on rehearsal of semantic information rather than visual perceptual information, perhaps because they needed to describe the pictures verbally in the free recall test, and that semantization and rehearsal were more present for high reward items. If this interpretation is correct, then although the direction of the effect is the opposite of that predicted, the reward-dependent decrease in ssVEP power across both mixed and pure lists may indeed act as a neural marker that matches eCMR’s predictions. Exactly how reward modulates ssVEPs may depend on the visual processing demands of the task. In two previous studies reward and subsequent memory increased ssVEP power, but these studies used motion detection (Grahek et al., 2021) and a recognition memory for visual shapes (Martens et al., 2012), both tasks which emphasized visual processing to a greater degree than here, where free recall of simple objects emphasized semantisation. While reward lowered ssVEP power here, it robustly increases ssVEP power in studies that employ negative emotional stimuli (Hajcak et al., 2013; Keil et al., 2008; Talmi, Slapkova, et al., 2019; Wieser et al., 2016), perhaps because such stimuli capture visual processing resources. At present, because the direction of the effect contradicts the pre-registered hypothesis, and its interpretation requires further research, we cannot consider the effect of reward on ssVEP markers of attention to offer strong evidence against the attentional borrowing account of list composition effects.

The ERP analysis showed that high reward items were related to more positive-going ERPs than low reward items, but only in mixed lists. The effect was widespread and included both centroparietal and frontal regions, so the spatial distribution overlaps with both an LPP and an FSW effect while it is more prolonged than a traditional P3 effect. The CSD analysis provided additional support for the interpretation that there were two separate ERP effects, one posterior (LPP) and one anterior (FSW) that overlapped in time. These late and prolonged ERPs are typically interpreted as indicative of strategic cognitive processing of stimuli. The ERP effects may reflect top-down selective attention to high reward items in mixed lists, as indexed by the more posterior LPP and/or selective rehearsal of high reward items in working memory, as indicated by the more anterior FSW. The similarity between the pattern of the ERP and behavioural results suggests that selective processing of high-reward items in mixed lists is likely to contribute to the reward list-composition memory effect, consistent with the selective encoding account and the attentional borrowing account. The finding that reward is related to an FSW contrasts with a recent study by Elliot et al. (2020) which did not observe an FSW effect during encoding in a value-based memory paradigm. That study used a recognition test which is known to be associated to less strategic encoding compared to free recall tests (Cohen et al., 2017), which may explain the discrepancy in results.

Reward-enhanced memory had previously been related to a posterior P3/LPP like effect, but reward manipulations are associated with a number of potential neural and cognitive processes, which previous work could not distinguish. The ERP results in our study shows that they are only present in mixed lists and is therefore context-dependent. This finding contradicts ERP studies of the emotion-based list-composition effect, which have observed an emotion-dependent increase in LPP amplitudes in both mixed and pure lists (Barnacle et al., 2018; Zarubin et al., 2020). The discrepancy suggests that list-composition effects for reward and emotion rely on at least partially different mechanisms. It is likely that encoding strategies are employed more in reward experiments than in emotion experiments, given that participants are instructed to try to remember high reward information in reward experiments, while there is no instruction to prioritise negative items in emotion experiments, and evidence that emotion attracts attention obligatorily (e.g. Pourtois et al. 2013; Talmi et al. 2007). Because emotional valence is intrinsic to the stimulus while reward needs to be associated with the stimulus, it is possible that top-down attention processes play a larger role in encoding of stimuli associated with reward. This interpretation agrees with the observation that the subsequent memory effect for emotional stimuli relies on regions associated with the bottom-up attention network, while the subsequent memory effect for neutral stimuli relied on the top-down attention network (Barnacle et al. 2016). The difference between the effect of reward and emotion may be best explained through the construct of ‘task usefulness’. Emotional items may be chronically useful because of an evolved sensitivity to threat-related content. By contrast, indication that certain items predict high reward items may only be ‘useful’ to participants in mixed lists. The effect of context on reward-dependent but not emotion-dependent modulations of the LPP may hint that this component is sensitive to ‘task usefulness’, rather than reward or emotion *per se*, mimicking recent results where reward did not enhance recall once usefulness was controlled for (Chakravarty et al. 2019).

The ssVEP and the LPP have both been suggested to reflect increased attention to emotional materials in studies of emotional stimuli. Prior research has shown that the two measures have different neural generators. In the context of emotion, the LPP has been suggested to be related to activity in a broad network including both cortical and subcortical regions (e.g. Keil et al., 2002; Liu et al., 2012; Sabatinelli et al., 2007, 2013) while flicker-induced ssVEPs mainly are generated in primary visual cortex and adjacent higher-order cortices (e.g. Russo et al., 2007; Wieser & Keil, 2011). However, this evidence cannot rule out that both LPP and ssVEP are different proximal manifestations of the same distal appraisal of a stimulus as motivationally salient; for example, both scalp measures could be due to downstream effects of amygdala activation (Pourtois et al., 2013). Measuring ssVEPs and LPP simultaneously makes it possible to investigate if these two measures capture the same aspect of attention. To the best of our knowledge, only one prior study has combined these two measures, but the authors used a very different orienting task of emotional attention, rather than the intentional encoding task in the present experiment (Hajcak et al., 2013). Nevertheless, even though Hajcak and colleagues found that negative emotion increased both the LPP effect and the ssVEP, the two measures were uncorrelated across participants, suggesting that they are measuring different aspects of attention to emotion. Our results provide further evidence for this possibility and extend it to effects of reward. Here, the two measures had a) opposite relation to reward (increased amplitude for the LPP in mixed lists and decreased power for the ssVEP) and b) differed in terms of the influence of list composition, where the LPP was list-context dependent while the ssVEP was not. Taken together, while both components may index increased attention, and may even be due to activations of the same driving influence, we show that they are functionally dissociated.

To conclude, the present study demonstrated that reward enhanced memory depends on list context. This result supports the prediction of eCMR as a computational model of reward-enhanced memory, but they also agree with selective encoding accounts. Going back to the question in the title of this article, our ERP results suggest that at least part of the context dependency of reward-enhanced memory is due to application of strategic processes to selectively encode high-reward items, but only in contexts in which they compete for cognitive resources with low-reward items. The selective encoding account is not a formal model, however, and does not make predictions for the dynamics of free recall. Because the dynamic pattern of recall matched the predictions of eCMR, our results suggest that the formal mechanisms this model embodies are useful to account for some of the effect of reward on free recall. Future computational work may allow retrieved-context models such as eCMR to account for strategic processes, something they are not presently able to simulate. Decreased ssVEP power in the high reward condition may be a neural marker of the tighter associations eCMR predicts link the context and the high-reward items, but we offer this interpretation cautiously, given the unpredicted direction of the effect, especially against previous literature and our own pre-registered hypothesis. Interestingly, the pattern of the LPP here is different from what have previously been observed in studies of list-context dependence in emotion and suggest that top-down encoding processes may be more important to explain list-composition effects for reward than for emotion. To test this hypothesis, future research could manipulate both emotion and reward, and examine whether the differences in their effects on the LPP become manifest even when they are measured in the same experiment. More broadly, the discussion highlights a number of putative differences between the influence of reward and emotion on the neural mechanism of intentional encoding. If corroborated, different formal mechanisms may be necessary to account for recalling different types of motivationally-significant experiences.

## Author note

We thank S Cradock for assistance in data collection and I. Laing for help in data coding. The study was supported by a joint award from the School of Biological Sciences, University of Cambridge, The Newton Trust, and Welcome Trust Institutional Strategic Support Fund (204845/Z/16/Z).

